# Inhibition of BK_Ca_ channels protects neonatal hearts against myocardial ischemia and reperfusion injury

**DOI:** 10.1101/2021.11.02.466585

**Authors:** Shridhar Sanghvi, Kalina Szteyn, Devasena Ponnalagu, Divya Sridharan, Alexender Lam, Inderjot Hansra, Ankur Chaudhury, Uddalak Majumdar, Andrew R. Kohut, Shubha Gururaja Rao, Mahmood Khan, Vidu Garg, Harpreet Singh

## Abstract

BK_Ca_ channels are large-conductance calcium and voltage-activated potassium channels that are heterogeneously expressed in a wide array of cells. Activation of BK_Ca_ channels present in mitochondria of adult ventricular cardiomyocytes is implicated in cardioprotection against ischemia-reperfusion (IR) injury. However, the BK_Ca_ channel’s activity has never been detected in the plasma membrane of adult ventricular cardiomyocytes. In this study, we report the presence of the BK_Ca_ channel in the plasma membrane and mitochondria of neonatal murine and rodent cardiomyocytes which protects the heart on inhibition but not activation. Furthermore, K^+^ currents measured in neonatal cardiomyocyte (NCM) was sensitive to iberiotoxin (IbTx), suggesting the presence of BK_Ca_ channels in the plasma membrane. Neonatal hearts subjected to IR when post-conditioned with NS1619 during reoxygenation increased the myocardial infarction whereas IbTx reduced the infarct size. In agreement, isolated NCM also presented increased apoptosis on treatment with NS1619 during hypoxia and reoxygenation, whereas IbTx reduced TUNEL positive cells. In NCMs, activation of BK_Ca_ channels increased the intracellular reactive oxygen species post HR injury. Electrophysiological characterization of NCMs indicated that NS1619 increased the beat period, field, and action potential duration, and decreased the conduction velocity and spike amplitude. In contrast, IbTx had no impact on the electrophysiological properties of NCMs. Taken together, our data established that inhibition of plasma membrane BK_Ca_ channels in the NCM protects neonatal heart/cardiomyocytes from IR injury. Furthermore, the functional disparity observed towards the cardioprotective activity of BK_Ca_ channels in adults compared to neonatal heart could be attributed to their differential localization.

## Introduction

Large conductance calcium and voltage-activated potassium channels (BK_Ca_ channels) encoded by the *Kcnma1* gene are the key electrochemical couplers between cellular metabolic state and Ca^2+^ homeostasis. The *Kcnma1* gene is highly conserved through a wide spectrum of species, including *Drosophila Melanogaster* [1, 2], *mus musculus* [3], and *homo sapiens* [4]. BK_Ca_ channels are expressed in a broad range of excitable and non-excitable cells and have been implicated in various fundamental physiological processes, including regulation of gene expression [5], urinary and erectile autonomic functions [6, 7], vascular tone [8], neuronal excitability [9], cardiac rhythmicity [10-12] and aging [2]. Recently, several *Kcnma1*-associated channelopathies have been identified in humans and BK_Ca_ channels have been projected as key therapeutic targets [4]. They are ubiquitously present in the plasma membrane of a majority of the cell types, and are highly selective for K^+^ with a large unitary conductance, and are sensitive to iberiotoxin (IbTx) as well as paxilline.

In addition to the plasma membrane, BK_Ca_ channels are also expressed in intracellular organelles like the endoplasmic reticulum, nuclei, lysosomes, and mitochondria [13, 14]. Channel expressed in the cell membrane and intracellular organelles is a product of the *Kcnma1* gene, but in cardiomyocyte mitochondria, the c-terminal splice variation of the *Kcnma1* gene (DEC) determines its inner mitochondria membrane (IMM) localization [13]. In physiological conditions, Ca^2+^ and voltage simultaneously control BK_Ca_ channel currents, but in the absence of Ca^2+^, membrane depolarization alone can generate BK_Ca_ channel currents. On the other hand, Ca^2+^ binding decreases the energy required to open the channel, shifting the open probability (P_O_) towards the more polarized state [15-17]. In smooth muscle cells, BK_Ca_ channels are key determinants in controlling the resting membrane potential and vascular tone, and hence, blocking the BK_Ca_ channel results in membrane depolarization and increased contractile tone [18-20]. BK_Ca_ channels are also present in the sinoatrial node (SAN) where their inhibition showed an increase in action potential duration (APD) [11]. Furthermore, when wild-type mice were injected with paxilline (a highly-specific BK_Ca_ channel inhibitor), their heart rate decreased by 30%, implicating BK_Ca_ channels in maintaining cardiac rhythmicity [12].

The use of pharmacological agents and genetically modified animal models demonstrated that the expression and opening of BK_Ca_ channels present in adult ventricular cardiomyocytes are directly involved in cardioprotection from IR injury [13, 21-24]. The cardioprotective mechanism is mediated by BK_Ca_ channels present in the mitochondria of adult cardiomyocytes [13, 21, 23]. On the other hand, there is growing evidence that BK_Ca_ channels are present in the plasma membrane at the embryonic stage in chick ventricular myocytes [25]. Potassium currents measured in the plasma membrane of cardiomyocytes from 10-12 days old chick embryos exhibited all the hallmarks of the BK_Ca_ channel properties such as: large single-channel conductance, voltage, and Ca^2+^ dependence and sensitivity to tetramethylammonium (TEA)- and charybdotoxin[25]. Additionally, human induced pluripotent stem cells-derived cardiomyocytes (hiPSC-CMs), showed sensitivity to IbTx treatment that shifted membrane polarization towards more positive potentials and caused a disturbance in action potential wave which was not observed in cardiomyocytes isolated from the adult human left ventricle [26]. However, the precise role and impact of BK_Ca_ channels in the plasma membrane of cardiomyocytes during the early stages of development are not yet elucidated.

Although activation of cardiomyocyte mitoBK_Ca_ channels has been long recognized in cardioprotection against ischemia-reperfusion injury in adults, the role and function of plasma membrane BK_Ca_ channels in neonates are completely unknown. Here, we present our novel findings showing a stage-dependent localization of BK_Ca_ channels in cardiomyocytes during early development, and its impact on cardiomyocyte function and cardioprotection.

## Materials and Methods

All experiments were conducted in accordance with guidelines and approved by The Ohio State University IACUC committee. C57BL/6NCrL mice and Sprague–Dawley (SD) were used. BK_Ca_^R207Q^ mice were obtained from Andrea Meredith (University of Maryland School of Medicine, Baltimore) and locally bred.

### Neonatal P3 cardiomyocytes isolation

Hearts were surgically excised from postnatal day3 (P3) pups (from rats and mice) and placed in a dissociation buffer (NaCl 16 mM, HEPES 20mM, Na_2_HPO_4_ 0.8mM, glucose 5.4 mM, KCl 5.4 mM, MgSO_4_ 0.8 mM, pH 7.35). Ventricles were minced in dissociation buffer followed by centrifugation at 200 x g for 5 min. at 4°C. The pellet was then resuspended in a dissociation buffer containing 0.25% Trypsin and the digestion was carried out at 37°C for 20 min in the water bath shaker. After 20 min, the cells were spun down (200 x g, 5 min, 4°C) and the dissociated neonatal cardiomyocytes were passed through a cell strainer (100 µm) and enriched with 20% (*v/v*) horse serum. The digestion of the remaining tissue pellet was repeated at least twice or until a clear tissue pellet was observed. The dissociated neonatal cardiomyocytes were then seeded on gelatin (0.1%, *w/v*) coated coverslips and cultured in DMEM containing 20% (*v/v*) FBS and penicillin (100 I.U.) / streptomycin (100 µg/ml), cultured in humidified 5% (*v/v*) CO_2_ incubators at 37°C and medium change every 2 days.

### Neonatal cardiomyocyte isolation (P7, P14, P21, and P28)

Cardiomyocytes were isolated from postnatal day 7 (P7), day 14 (P14), day 21 (P21), and day 28 (P28) pups, respectively using the simplified Langendorff-free method of isolation (1). Briefly, pups were anesthetized, and the chest was cut open to expose the heart. The descending aorta was cut followed by injection of 7 ml EDTA buffer [NaCl 130 mM, KCl 5 mM, NaH_2_PO_4_ 0.5 mM, HEPES 10 mM, glucose 10 mM, Butanedione Monoxime (BDM) 10 mM, Taurine 10 mM, EDTA 5 mM, pH 7.8] into the right ventricle. The ascending aorta was clamped, and the heart was transferred to a Petri dish containing EDTA buffer. The left ventricle was injected with 10 ml of EDTA buffer followed by 3 ml of perfusion buffer (NaCl 130 mM, KCl 5 mM, NaH_2_PO_4_ 0.5 mM, HEPES 10 mM, glucose 10 mM, BDM 10 mM, Taurine 10 mM, MgCl_2_ 1mM, pH 7.8), and 10-20 ml of collagenase enzyme solution (collagenase type 2 0.5 mg/ml, collagenase type 4 0.5 mg/ml, protease XIV 0.05 mg/ml prepared in perfusion buffer). The heart chambers were separated and gently pulled into 1-mm pieces. The cells were dissociated using gentle trituration and the collagenase activity was inhibited by 5 ml of stop buffer [perfusion buffer containing 5% (*v/v*) sterile fetal bovine serum (FBS)]. The cells were washed three times with perfusion buffer and allowed to settle with gravity.

### Adult cardiomyocyte isolation

Animals were injected intraperitoneally with heparin (200 IU/kg) and 20 min later they were anesthetized in a 4% isoflurane chamber. Hearts were then harvested and instantaneously arrested in ice-cold PBS (KCl 2 mM, KH_2_PO_4_ 1.5 mM, NaCl 138 mM, Na_2_HPO_4_ 8.1 mM) to remove excess blood. Hearts were transferred to ice-cold Tyrode’s solution (NaCl 130 mM, KCl 5.4 mM, MgCl_2_ 1 mM, Na_2_HPO_4_ 0.6 mM, Glucose 10 mM, Taurine 5 mM, BDM 10 mM, and HEPES 10 mM, pH 7.4, oxygenated with 95% (*v/v*) O_2_-5% (*v/v*) CO_2_], and mounted on a modified Langendorff apparatus at a constant pressure of ∼5-7 ml/min. After 5 min of perfusion at 37 °C with Tyrode’s solution, the mice hearts were perfused for 12-15 min with Tyrode’s solution containing 186 U/mL Collagenase Type-2 and 0.5 U/mL Protease Type-XIV, and then washed for 5 min with a high Potassium buffer (KB) [KCl 25 mM, KH_2_PO_4_ 10 mM, MgSO_4_ 2 mM, Glucose 20 mM, Taurine 20 mM, Creatine 5 mM, K-Glutamate 100 mM, Aspartic acid 10 mM, EGTA 0.5 mM, HEPES 5 mM, and BSA 1% (*w/v*), pH 7.2-7.3 oxygenated with 95% O_2_-5% (*v/v*) CO_2_]. For rat hearts, the enzyme-Tyrode solution was perfused for ∼20 min and contained 372 U/mL Collagenase Type-2 and 1.0 U/mL Protease Type-XIV, while washing with KB was for 15 min. After washing, the left ventricle was cut into pieces in KB to dissociate cells. Isolated ventriculocytes were filtered (100 μm strainer), and gravity settled for 10 min on ice.

### Hypoxia/Reoxygenation treatment

Neonatal cardiomyocytes were subjected to hypoxia/reoxygenation injury. After 4 days of culturing the neonatal cardiomyocytes with 70% confluency in cell culture dishes or coverslips coated with 0.1% gelatin, cells were subjected to hypoxia in a modular humidified 37°C hypoxia chamber (Biospherix, C127) with 1% O2, 5% CO2 and balanced N_2_ for 6 hr. After hypoxia incubation, cells were conditioned with DMSO control, BK_Ca_ channel inhibitor (iberiotoxin), and activator (NS1619) for 30 min followed by reperfusion with DMEM supplemented with 10%FBS and 1% Penicillin-Streptomycin antibiotics.

### Total RNA isolation and qPCR

Total RNA from the gravity-settled cardiomyocytes was then purified using TRIzol reagent (Invitrogen) followed by digestion with on column-RNase-free DNAse digestion kit (Qiagen) and clean up with RNeasy mini kit (Qiagen). Cleaned-up RNA (0.5 µg) was reverse transcribed with Omniscript Reverse Transcription kit (Qiagen) using oligo dT primers in a 20 µL reaction volume. Real-time PCR was performed using SYBRTM Green master mix (Applied Biosystems), 1 µL of RT reaction product, and 200 nM primer pairs (**supplementary table 1**) in a 10 µL reaction volume. The thermal cycling conditions included an initial denaturation at 95°C for 5 min, and 40 cycles of 95°C for 45 s, 60°C for 45 s, and 72°C for 45 s. The controls used in the qPCR are (-)RT (cDNA with no reverse transcriptase), Water control (water instead of the template), and primers used to amplify GAPDH (2). All samples were run in duplicates. The efficiency of primers was calculated to be more than 94% for BK_Ca_-DEC and BK_Ca_-Total (3). The fold change in expression of BK_Ca_-Total and BK_Ca_-DEC have been plotted relative to GAPDH.

### Western Blot Analysis

Cardiomyocytes isolated from wild type and *Kcnma1*^*-/-*^ mice were lysed with modified RIPA buffer (Tris-HCl 50 mM, NaCl 150 mM, EDTA-Na_2_ 1 mM, EGTA-Na_4_ 1 mM, Na_3_VO_4_ 1 mM, NaF 1 mM, Nonidet P-40 1% (v/v) Na-deoxycholate 0.5% (wt/vol), and SDS 0.1% (wt/vol), pH 7.4) containing protease inhibitors (1 tablet/50 mL; Roche) and PhosSTOP(tm) (1 tablet/10 mL; Roche), flash freeze in liquid nitrogen and incubated 1 hr at 4°C. Samples were centrifuged for 20 min at 10,000 × g and the supernatants were collected as lysate. Proteins (50 μg/lane) were separated on 4-20% SDS/PAGE and transferred to nitrocellulose membranes. Loading was corroborated with Ponceau S staining. Nitrocellulose membranes were blocked with Intercept® blocking buffer at room temperature for 1 hr and incubated overnight with anti-BK_Ca_ pAb (2 μg/mL, Alomone labs, APC21) and anti-Dynamin I mAb (2 μg/mL). Membranes were washed thrice with 1X Tris-buffered Saline containing Tween-20 and incubated with 0.01 μg/mL secondary Abs (IR-dye 800 goat anti-rabbit IgG and IR-dye 680 goat anti-mouse) for 60 min at room temperature. After extensive washing, membranes were visualized using BioRad ChemiDoc MP.

### Immunocytochemistry

Rodent and mouse neonatal cardiomyocytes were incubated with wheat germ agglutinin (WGA) at 37°C on ice for 60 min and/or mitotracker for 10 min at 37°C. Cells were fixed with 4% (*w/v*) PFA and permeabilized with 0.5% (*v/v*) Triton-X-100. Neonatal cardiomyocytes were incubated with anti-ATP synthase (mouse, Abcam), anti-BK_Ca_ antibodies, and cytochrome C (CST) overnight at 4°C. Secondary antibodies conjugated with anti-mouse Alexa-488 and anti-rabbit Atto-647N were added for 60 min at room temperature. To label nuclei, DAPI was added (1:10,000) in the wash solution. Coverslips were mounted with Mowiol®. Cells were imaged with Nikon A1R high-resolution confocal microscopy. The colocalization index was calculated using protein proximity index analysis (4). Images were filtered by custom-built software as described earlier.

### TUNEL assay

The extent of the apoptotic damage was monitored using a deoxynucleotidyl transferase dUTP nick end labeling (TUNEL) assay kit (Thermo Scientific; C10618). Briefly, P3 neonatal cardiomyocytes were cultured on 24-well plates coated with 0.1% gelatin at 70% confluency for 96hr, hypoxia injury, and postconditioning with DMSO, NS1619, IbTx, and Pax was induced as described earlier. After 12 hr of reoxygenation, the cells were fixed with 4% (*v/v*) PFA for 10 min, permeabilized with 0.25% (*v/v*) Triton-X-100 for 20 mins, and apoptotic nuclei were stained using a TUNEL assay kit as described by the manufacturer’s protocol. Fluorescent images were taken using Nikon A1R confocal microscope.

### Cellular ROS measurement

Reactive oxygen species production was measured using CM-H2DCFDA (Life Technologies, C26827). Neonatal cardiomyocytes cultured on coverslips subjected to 6hr hypoxia injury, conditioned with DMSO, Iberiotoxin (100 nM), or NS1619 (10 μM) for 30 min following 12hr reperfusion injury. Cells were stained with 10 μM of CM-H2DCFDA in regular DMEM media without phenol red for 30 mins. Cells supplemented with 10% (*v/v*) FBS and 1% (*w/v*) PenStrep. Images were acquired using Nikon A1R confocal microscope.

### Patch-clamp

Patch-clamp experiments were performed on mouse and rat neonatal cardiomyocytes 96hr post isolation. Cells were cultures on coverslips coated with poly-D-lysine. Patch electrodes were fabricated from borosilicate glass (Sutter Instrument, Navato, CA) on an M-87 horizontal puller (Sutter Instrument, Navato, CA), with an average resistance of 3-6 MΩ. All the recordings were performed at room temperature and currents were acquired using an EPC10 USB amplifier (HEKA Electronic, Germany) with accompanying PatchMaster (HEKA, Germany) software that was also used for analysis. Patch-clamp experiments were performed in whole-cell and inside-out mode. The bath solution contained (KCH_3_SO_3_ 140 mM, MgCl_2_ 2 mM, KCl _2_ mM, HEPES 20 mM, pH 7.3). Pipette solution contained (KCH_3_SO_3_ 140 mM, CaCl_2_ 10 mM, HEDTA 5 mM, HEPES 20 mM, pH 7.3. During whole-cell experiments cells were held at -70 mV, 40 ms pulses were applied from -70 to +150 mV in 20 mV steps. After establishing stable baseline currents 100 nM IbTx was added to the bath solution, to validate the ion channel responsible for detected K^+^ currents.

### Langendorff isolated perfused neonatal rat hearts

Six-day-old Sprague Dawley (SD) rat pups were injected intraperitoneal with heparin (100 IU/Kg) to prevent blood coagulation. After 20 min, animals were anesthetized in an isoflurane chamber. Hearts were rapidly excised and arrested in ice-cold filtered Krebs-Henseleit (KH) buffer containing (Glucose 11.1 mM, NaCl 118 mM, KCl 4.7 mM, MgSO_4_ 1.2 mM, KH_2_PO_4_ 1.2 mM, NaHCO_3_ 25 mM, and CaCl_2_ 2 mM at pH 7.4). The hearts were cleaned to remove non-cardiac tissue and excess fat before cannulating the aorta onto a 21-gauge cannula. The hearts were retrogradely perfused with KH buffer at a constant rate of 2-3 ml/min and maintained at 37°C in the water-jacketed Langendorff chamber. The buffer was constantly aerated with a mixture of 95% O_2_ (*v/v*) and 5% CO_2_ (*v/v*). After equilibrating the hearts by perfusing with KH buffer for 15 min, the hearts were subjected to global normothermic ischemia for 20 min by stopping the perfusion. Following the ischemia, hearts were then post-conditioned with NS1619 (10 μM), IbTx (100 nM), and DMSO (0.001%) for 5 min and reperfused with KH buffer for 60 min at 37°C.

### Myocardium infarct size

Myocardium infarction was measured by staining the heart sections with 2, 3, 5-triphenyl tetrazolium chloride (TTC) stain. After reperfusion for 60 min, the hearts were cut into five transverse sections parallel to the atrioventricular groove. Heart slices were incubated in 1% TTC (w/v) at 37°C for 20 min followed by fixation in 4% PFA (*w/v*). The stained sections were imaged using Leica S9i. To demarcate the infarcted region (pale) versus viable myocardial tissue (brick red) ImageJ was used to quantify the area. The total infarcted area was calculated against the total heart sections and expressed as the percentage.

### MEA measurements

Rat neonatal cardiomyocytes were isolated from P3 pups using the Pierce primary cardiomyocyte isolation kit (Thermo Scientific). The neonatal cardiomyocytes were seeded on (50 μg/ml) fibronectin-coated CytoView MEA-24 well plates (Axion BioSystems, Inc.) at 100,000 cells per well and data were acquired 72 hr post isolation using Maestro Edge MEA platform (Axion BioSyetems, Inc.). The voltage data were sampled for 16 electrodes/well simultaneously at (15 µV). LEAP inductions were performed using the AxIS Navigator software (Axion BioSystems, Inc.) using the dedicated stimulator in the Maestro Edge on selected planar microelectrodes on 24 well CytoView plates. For data analysis of (1) cardiac field potential duration (FPD), (2) beat period, (3) spike amplitude, and (4) conduction velocity were plotted as fold change after comparing with treatment groups (DMSO, IbTx, or NS1619) to their respective baseline wells. Graphical representation of the data was automatically detected and plotted using CiPA Analysis Tool software (Axion BioSystems, Inc.). Cardiac LEAP signals were detected using AxIS Navigator and representative data shown were plotted against DMSO control. Cardiac LEAP signals (a) rise time and (b) action potential duration (APD), APD30, APD50, and APD90 against DMSO were calculated and detected using CiPA Analysis Tool.

### Immunohistochemistry of human hearts

Human right ventricular myocardial sections were obtained from the Heart Center Biorepository at Nationwide Children’s Hospital. This study was approved by the Institutional Review Board (IRB) at Nationwide Children’s Hospital (IRB #07-00298). The myocardial sections were from infants under 3 months of age, with the following congenital heart defects (tetralogy of Fallot, truncus arteriosus, and pulmonary stenosis). The myocardial tissues were fixed in PFA and sectioned for histological analysis. For immunohistochemical staining, tissue sections were deparaffinized in xylene and rehydrated in grades of ethanol and PBS, followed by antigen retrieval using citrate-based buffer (Vector Laboratories# H-3300-250). Sections were incubated at room temperature with 3% (*v/v*) H_2_O_2_ for 10 min to quench endogenous peroxidase activity and blocked by 5% (*v/v*) normal goat serum (Vector Laboratories, #S-1000) in PBS for 1 hour. After blocking, sections were incubated with WGA (1:500) for 60 min at room temperature to the label plasma membrane. After 60 minutes sections were washed with PBS three times. Sections were incubated in 0.5% (*v/v*) PBS-Triton-X-100 for 10 minutes at room temperature for permeabilization. After washes with PBS sections were incubated in a primary antibody against BK_Ca_ channels (Alomone Labs, APC021) overnight at 4°C. Following primary antibody incubation, sections were incubated with secondary antibodies. During washing DAPI (1:10,000 dilution) was added and sections were washed with PBS. Cardiac sections were visualized with a high-resolution confocal microscope (Nikon A1R).

### Statistical analysis

Data were analyzed using Prism (GraphPad) or Excel and reported as mean ± SEM. To assess the significant difference, comparisons were measured between two groups using a paired one-tailed Student’s t-test. The significance between the groups in apoptotic nuclei and ROS was compared using a one-tailed unpaired student’s t-test followed by a non-parametric Mann-Whitney test for comparison between the ranks. *P* ≥ 0.05 indicated non-significant, *P* ≤ 0.05 indicated significant.

## Results

### BK_Ca_ channels are present in the plasma membrane of neonatal cardiomyocytes

BK_Ca_ channels are known to localize to the mitochondria of adult mouse and rat ventricular cardiomyocytes [13, 22, 23]. However, their localization in NCMs is unknown. We evaluated the the localization of BK_Ca_ channel in NCMs isolated from rats and mice using a highly-specific anti-BK_Ca_ antibody [13, 22] (**Fig. 1 E**). Similar to adult ventricular cardiomyocytes [13, 22], in both rat and mouse NCMs, BK_Ca_ channels showed a strong localization to mitochondria (co-labeled with anti-ATP5A) (**Fig. 1**). In rats, localization of BK_Ca_ channels to mitochondria increased from 37 ± 5% (n=20) in postnatal day 3 (P3) NCMs to 56 ± 4% (n=20) in adult ventricular cardiomyocytes. Similarly in mice, BK_Ca_ channels localization to ATP5A labeled-mitochondria was 40 ± 3% (n=20) which increased to 62 ± 4% (n=25) in P3 NCMs and adult ventricular cardiomyocytes, respectively. These findings were similar to earlier reports on the preferential localization of BK_Ca_ channels to cardiomyocyte mitochondria [27]. In adult ventricular cardiomyocytes isolated from mice or rats, there was a small amount of localization of BK_Ca_ channels to the plasma membrane 12 ± 1% (shown by WGA staining) which could be attributed to mitochondria present near the plasma membrane (**Fig. 1B, D, and F**). However, in NCMs isolated from rats and mice, BK_Ca_ channels showed 26 ± 3% (n=20) and 32 ± 3% (n=15), localization to WGA labeled plasma membrane (**Fig 1A, C, and F**), respectively.

**Figure 1:**
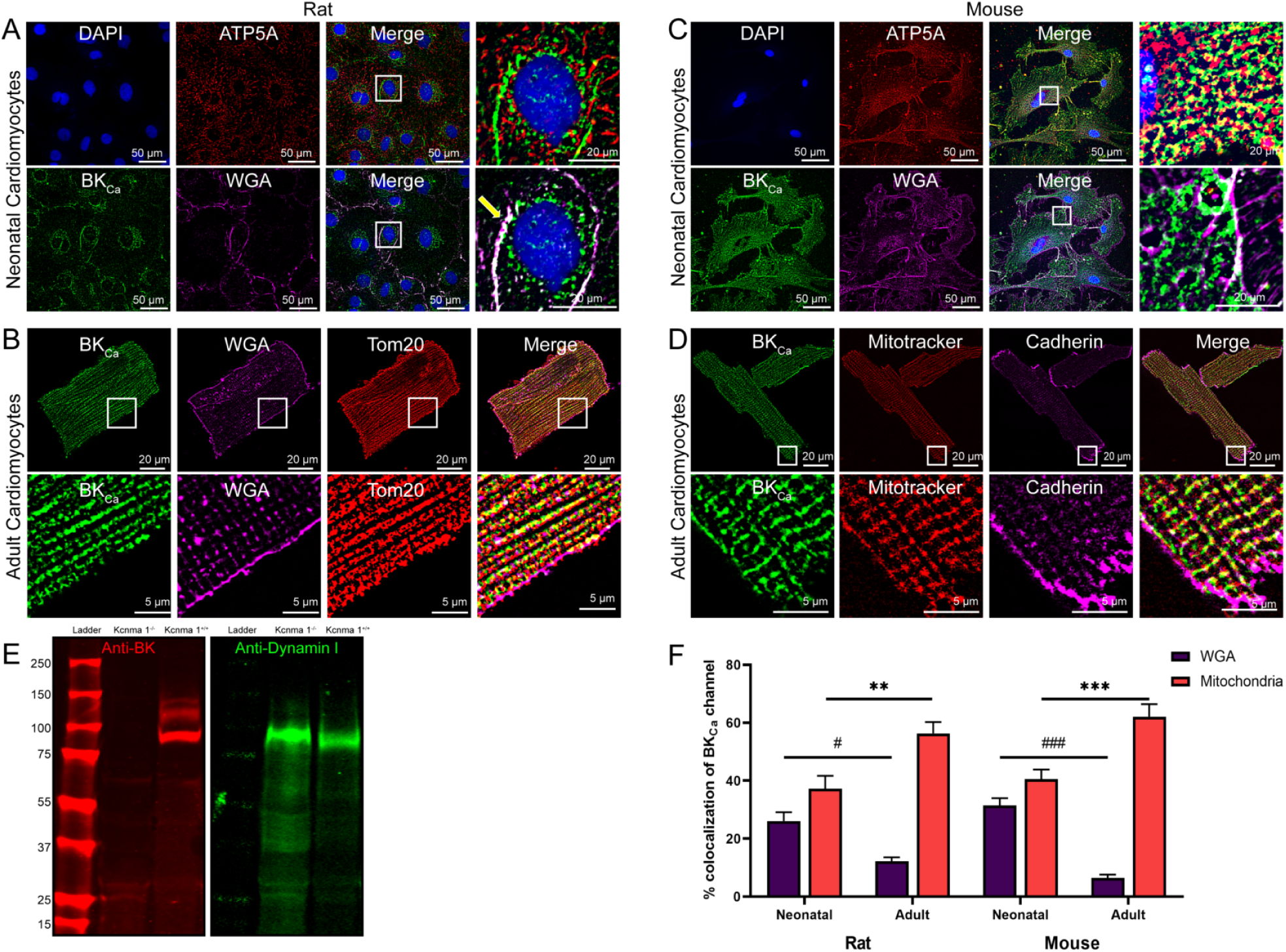
Localization of native BK_Ca_ channel in neonatal and cardiomyocytes isolated from the rodent heart. (A, B, C, and D) Isolated cardiomyocytes loaded with WGA (magenta) or MitoTracker (red), fixed, permeabilized, and labeled with BK_Ca_ (green), ATP5A, TOM20 (red), and Cadherin (magenta) antibodies. Nuclei were stained with DAPI (blue). Scale bars are 50 µM. The right panels in A, B, C, and D show merged images at higher magnification. (E) Validation of BK_Ca_ antibody in cardiomyocytes isolated from adult *Kcnma1*^*+/+*^ and *Kcnma1*^*-/-*^ mice. Dynamin 1 is used as a loading control. Molecular weights are given in KDa. (F) The purple bar represents percentage colocalization between the BK_Ca_ channel and plasma membrane (WGA) which is significantly higher in neonatal cardiomyocytes compared to adult cardiomyocytes. The red bar represents percentage colocalization between the BK_Ca_ channel and mitochondria which is lower in neonatal cardiomyocytes as compared to adult cardiomyocytes isolated from rodent hearts. All experiments were repeated independently at least four times and colocalization data are represented as mean ± SEM from 10 cells in each ‘n’ number. *P* values were determined by a one-tailed paired student’s t-test; ** ≤ 0.001, *** ≤ 0.0001, # ≤ 0.000001, ### ≤ 1.0*10^−12^.

### BK_Ca_ channels are functional in the plasma membrane of neonatal cardiomyocytes

Previous studies [14, 28, 29] have shown that BK_Ca_ channel-specific currents are not present in the membrane of adult ventricular cardiomyocytes, due to its absence in the plasma membrane [13]. Since we observed localization BK_Ca_ channels in the plasma membrane of NCMs, we evaluated whether these BK_Ca_ channels are functional NCMs using patch-clamp analysis. In the whole-cell configuration, cells were held at -70 mV holding potential and voltage steps of 20 mV (40 ms) from -70 to +150 mV were applied. As shown in **Fig. 2 A** and **D**., we recorded a large outward current in NCMs isolated from both rats and mice. In rat NCMs, the open probability of potassium currents was decreased by 46 ± 5% (n=10, **Fig. 2A, B, and C**) and in mouse NCMs, 32 ± 5% (n=10, **Fig.2 D, E**, and **F**) after application of IbTx, indicating the presence of functional BK_Ca_ channels in the plasma membrane of NCMs.

**Figure 2:**
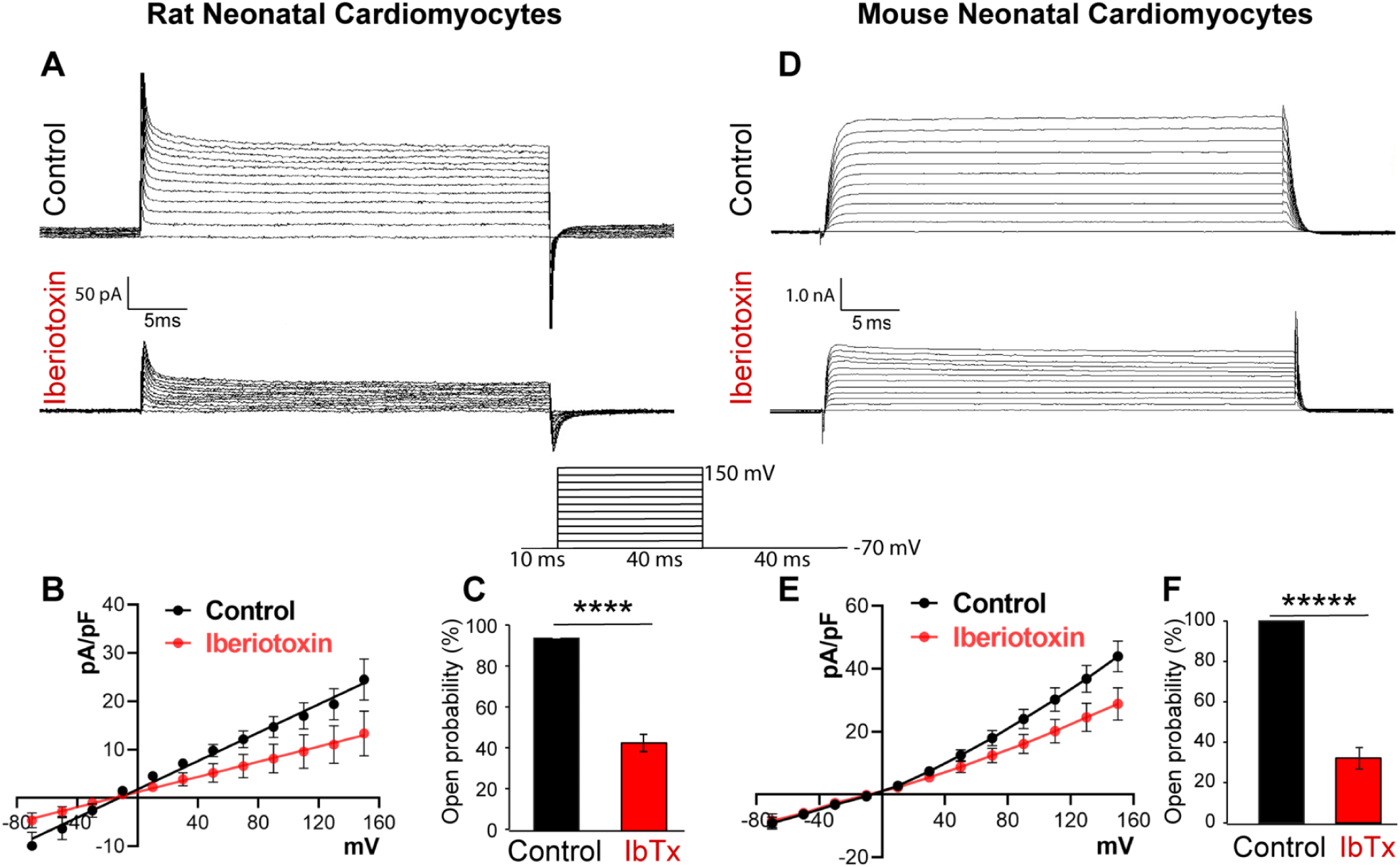
Rodent neonatal cardiomyocytes express functional BK_Ca_ channels. Traces of whole-cell BK_Ca_ channel currents were recorded from (A) rat neonatal cardiomyocytes and (D) mouse neonatal cardiomyocytes with or without iberiotoxin (100 nM). Neonatal cardiomyocytes (B) rat and (E) mouse, membrane currents were activated by 40 ms voltage steps of 20 mV between -70 mV to +150 mV from resting -70 mV holding potential. The current *versus* voltage relationships are plotted with or without iberiotoxin. (C and F) Percentage block of BK_Ca_ current in presence or absence of iberiotoxin. Cells were isolated from ten independent litters from mice and rats, and data are represented as mean ± SEM. *P* values were determined by a one-tailed paired student’s t-test; **** ≤ 0.00001, ***** ≤ 0.000001.

### Localization of BK_Ca_ channel during cardiomyocyte development

Due to the changes in the localization of BK_Ca_ channels from the plasma membrane to mitochondria in neonatal and adult cardiomyocytes, we investigated changes in their age-specific localization. Hence, we studied the localization of BK_Ca_ channels in cardiomyocytes isolated from pups at P3, P7, P14, P21, P28 days, and adult rats. In NCMs, BK_Ca_ channels were present in the plasma membrane as well as in mitochondria (**Fig. 3A-F**). Protein proximity index analysis [30] showed that BK_Ca_ localization to mitochondria increases from P3 to the adult stage, while localization of BK_Ca_ to plasma membrane decreases (**Fig. 3G**). The major determining factor attributed to the localization of BK_Ca_ to mitochondria is the C-terminus DEC splice variant (BK_Ca_ DEC) [13]. We quantified the DEC splice variant and compared it with the total BK_Ca_ (BK_Ca_ FL) signal present in rat cardiomyocytes isolated at different ages. We observed an increased expression of the BK_Ca_ DEC from P3 to adult which corresponds to increased localization of BK_Ca_ channels to mitochondria in cardiomyocytes **(Fig. 3G and H)**.

**Figure 3:**
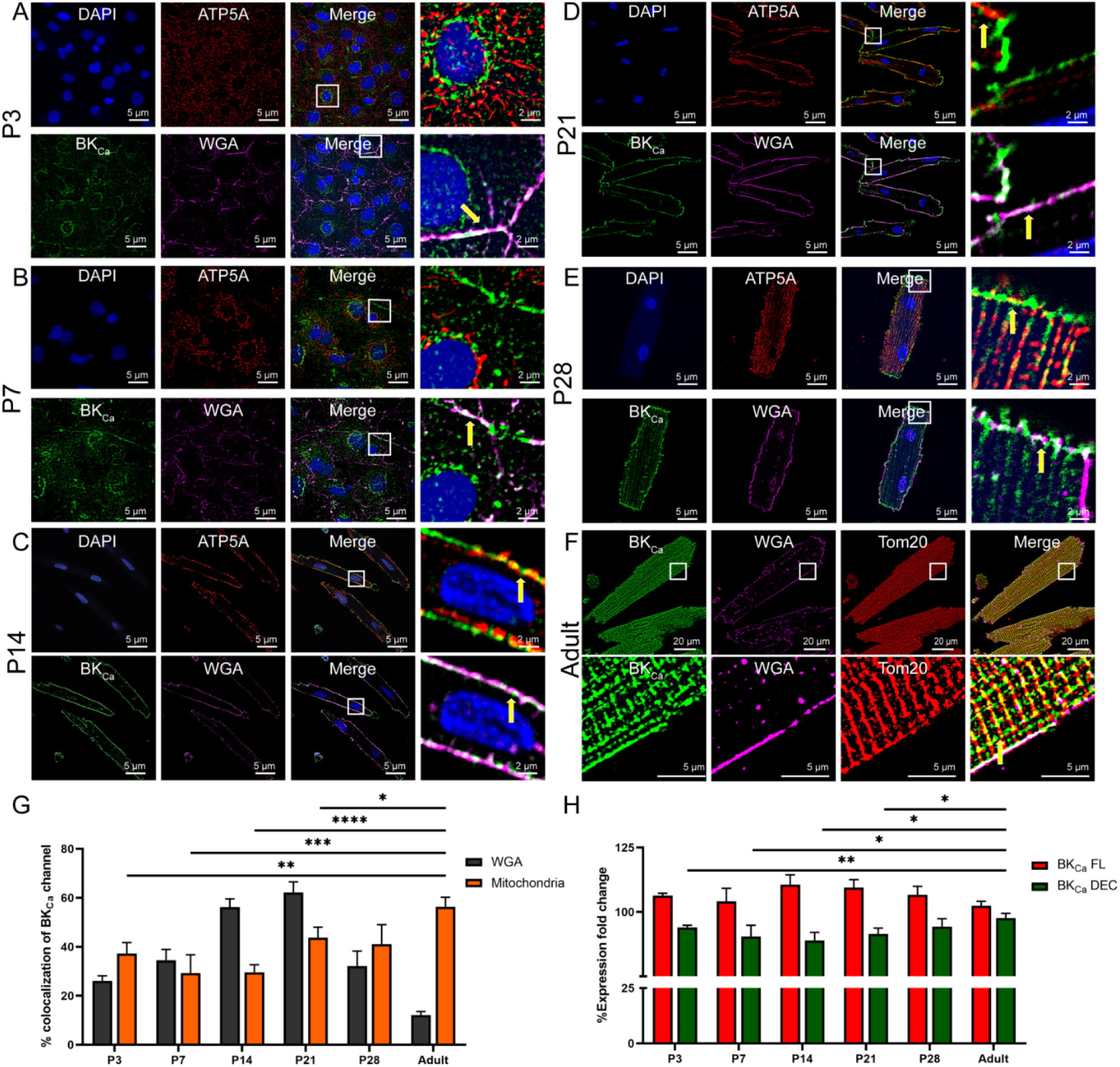
Mitochondrial targeting BK_DEC_ splicing increases with age in neonatal cardiomyocytes. (A-F) Cardiomyocytes isolated from aging (P3, P7, P14, P21, P28, and adult) rats loaded with WGA (Magenta), fixed, permeabilized, and labeled with BK_Ca_ (Green) and ATP5A (Red). The nucleus was stained with DAPI (blue). The right panels of A-F show colocalization of merged images between BK_Ca_ to WGA and BK_Ca_ to ATP5A at higher magnification. (G) Protein proximity index analysis showing percentage colocalization of BK_Ca_ to WGA decreases whereas BK_Ca_ to mitochondria increases in cardiomyocytes isolated from P3 to the adult stage. (H) Relative levels of BK_Ca_ FL and BK_Ca_ DEC mRNA expression in cardiomyocytes isolated from P3, P7, P14, P21, P28, and adult hearts were measured by RT-qPCR and were normalized to their relative GAPDH expression. Data represented as fold change to P3 BK_Ca_ FL and P3 BK_Ca_ DEC; mean ± SEM; at least 3 independent experiments. *P* values were determined by a one-tailed paired student’s t-test; * ≤ 0.05, ** ≤ 0.001, *** ≤ 0.0001, **** ≤ 0.00001.

### Activation of BK_Ca_ channels in neonatal rat heart increases myocardial infarction

Activation of BK_Ca_ channels either pharmacologically [13, 23, 24, 31-33] or genetically [22] in adult animal models results in cardioprotection from IR injury whereas inhibition of BK_Ca_ channels increases myocardial infarction[21]. We tested whether activation of BK_Ca_ channels can protect neonatal hearts from IR injury as observed in adult hearts. Hearts isolated from 6 days old rats (P6) were subjected to ischemia and immediate post-conditioning with a BK_Ca_ channel activator (10 µM NS1619) or inhibitors (100 nM IbTx) (**Fig. 4A**). Hearts were reperfused after post-conditioning. Infarct size was measured by staining cardiac sections at the end of reperfusion by 2, 3, 5 -triphenyl tetrazolium chloride (TTC) (**Fig. 4B-D**). Post-conditioning with NS1619 significantly increased the infarct size (58±3%) as compared to vehicle control (48 ± 5%) as shown in **Fig 4B, D**, and **E**. In contrast, hearts post-conditioned with IbTx showed reduced infarct size (32 ± 5%) (**Fig. 4C** and **E**) compared to the vehicle. To corroborate the expression of functional BK_Ca_ channels on the plasma membrane, we recorded potassium currents in NCM isolated from P6 rat pups. The open probability of potassium currents was significantly reduced by 33 ± 3% after the application of IbTx **(Fig. 4F)**. Thus our results indicate that contrary to adults, in neonates, the activation of the BK_Ca_ channel after IR injury is cardio-deleterious whereas BK_Ca_ channel inhibition is cardioprotective.

**Figure 4:**
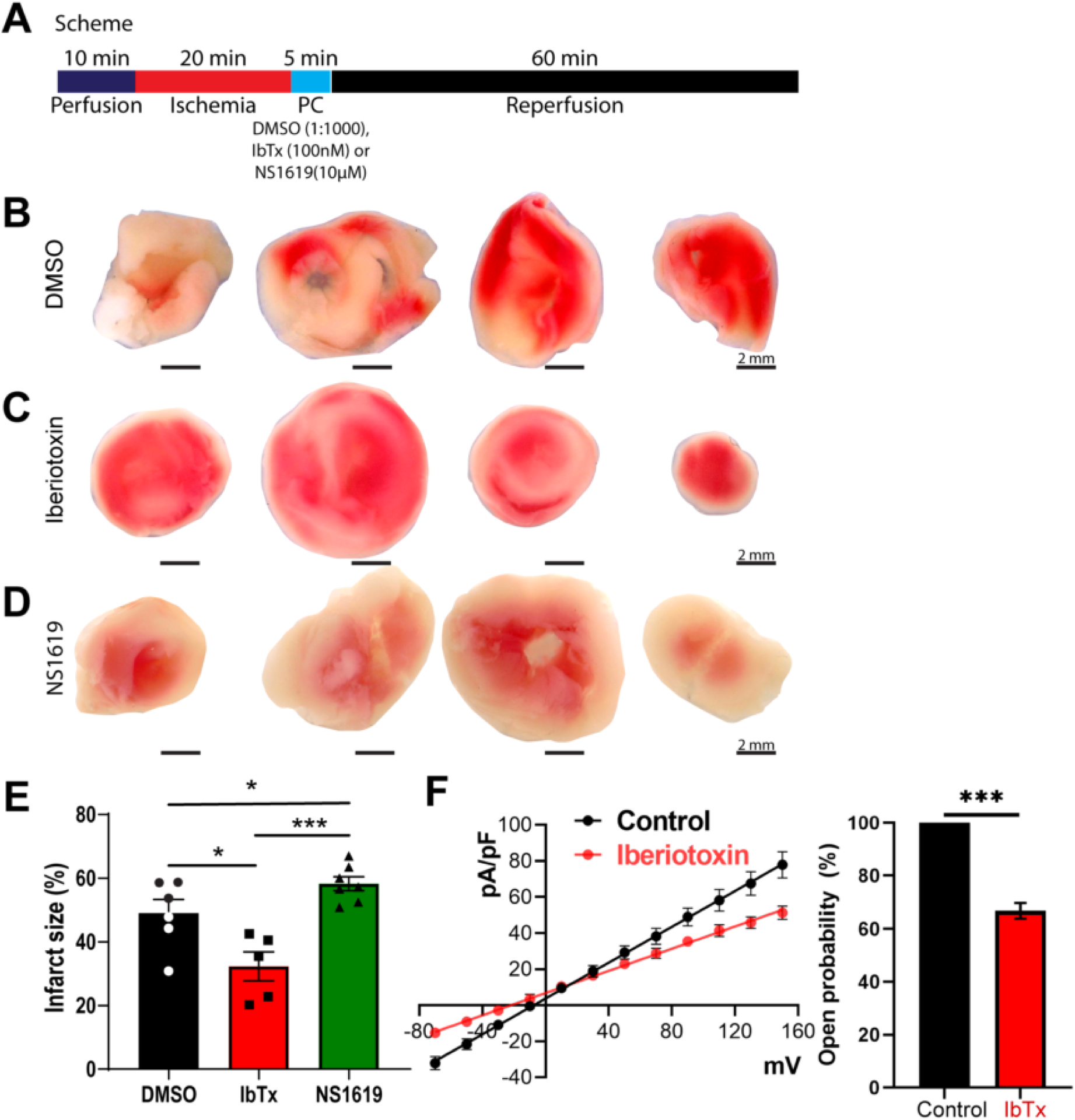
Inhibition of BK_Ca_ channel protects the neonatal heart from ischemia injury. (A) Schematics of the ischemia-reperfusion protocol. Rat P6 pup hearts were subjected to 20 min ischemia, 5 min post-conditioned with DMSO, Iberiotoxin (100 nM), or NS1619 (10μM) followed by 60 min of reperfusion. Hearts post-conditioned with (C and E) iberiotoxin exhibited significantly less infarction (white) as compared to the (B and E) DMSO control. In hearts post-conditioned with NS1619 displayed significantly higher infarction compared to DMSO control and IbTx treatment (E). The current versus voltage relationships are plotted with or without iberiotoxin (F). Percentage block of BK_Ca_ current in presence or absence of iberiotoxin. Data represented as mean ± SEM for percentage infarction from four independent experiments and electrophysiology data represented from four independent litters P6 rat pups. *P* values were determined by a one-tailed paired student’s t-test; * ≤ 0.05, *** ≤ 0.0001.

### Activation of the BK_Ca_ channel induces apoptosis after hypoxia-reoxygenation in NCM

One of the major roles attributed to the activation of BK_Ca_ channels is protecting adult cardiac cells and tissues from ischemia/hypoxia-reoxygenation injury[13, 21, 23, 34]. These functions are specifically ascribed to the presence of BK_Ca_ channels located in mitochondria of adult ventricular cardiomyocytes [21, 34]. Since we discovered BK_Ca_ channels in the plasma membrane of NCMs **(Fig. 1** and **2**), we determined the effect of plasma membrane BK_Ca_ channels in isolated NCMs after hypoxia-reoxygenation injury. We used cell impermeable (IbTx, 100 nM) to tease out the role of plasma membrane BK_Ca_ channels in HR-induced apoptosis in NCMs (P3). IbTx showed comparable apoptotic positive nuclei 21 ± 8% after HR injury to DMSO control 30 ± 9% as indicated by TUNEL assay (**Fig. 5A, B** and quantified in **D**). Surprisingly, NS1619 treatment showed 63 ± 14% TUNEL positive cells as compared to DMSO control 30 ± 8% after HR injury (**Fig. 5 A. C** and **D**).

**Figure 5:**
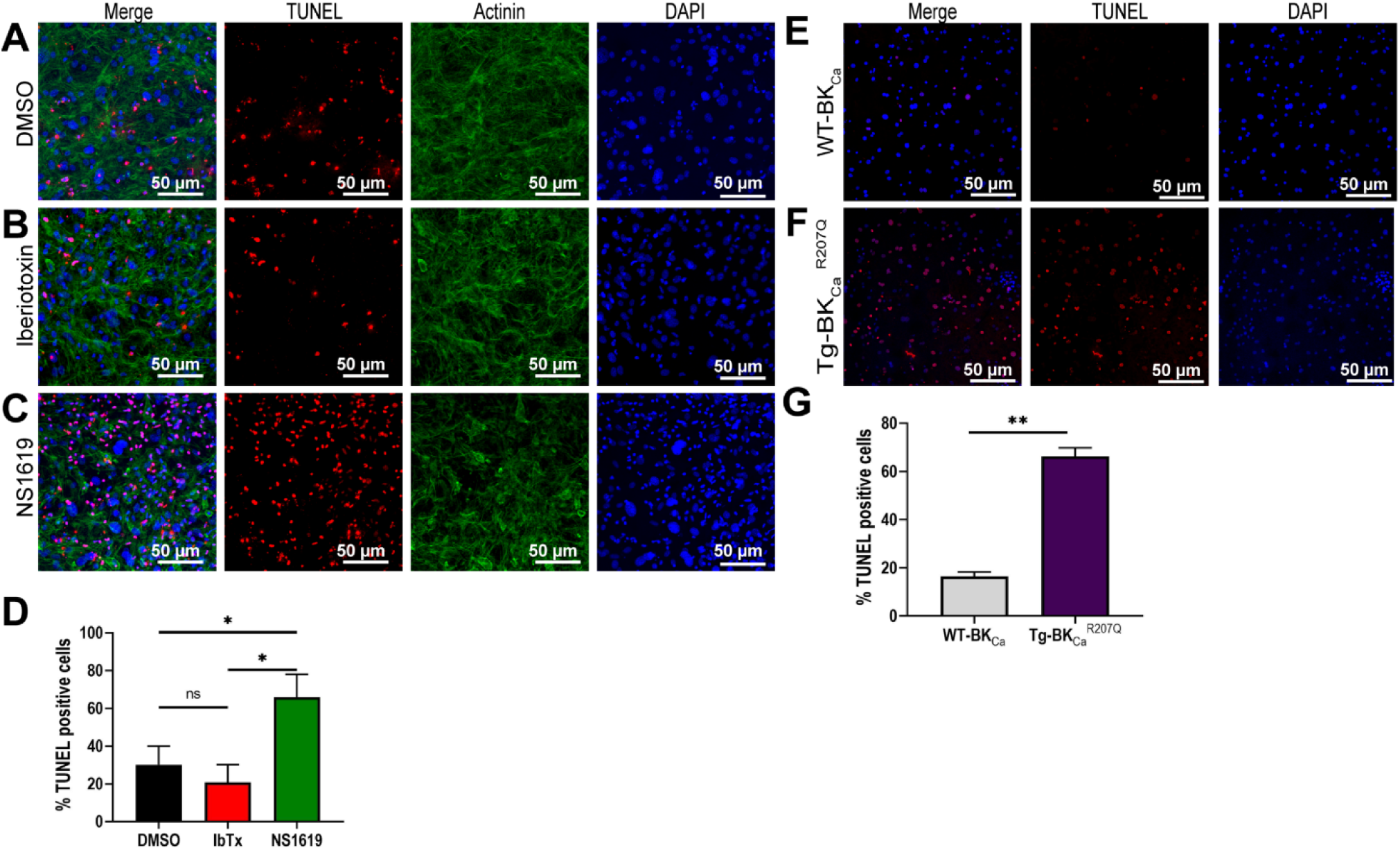
Inhibition of BK_Ca_ channel protects the neonatal cardiomyocytes from apoptotic damage post hypoxia injury. Isolated mouse neonatal cardiomyocytes were subjected to 6 hr hypoxia at 1% O_2_, 5 mins of post-conditioning with DMSO, Iberiotoxin (100 nM), or NS1619 (10μM) followed by 12 hr of reoxygenation at 21% O_2_. Cells were fixed and stained for TUNEL positive nuclei (red), Actinin (green), and Nucleus (blue). A total of 100,000 cells were seeded per assay. (B and D) Cardiomyocytes post-conditioned with IbTx shows fewer TUNEL positive nuclei compared to DMSO control (A and D). Cardiomyocytes post-conditioned with NS1619 exhibited a significantly higher number of TUNEL positive nuclei compared to DMSO and IbTx (D). Cardiomyocytes isolated from (E) wild-type pups (WT-BK_Ca_) or pups expressing (F) genetically activated BK_Ca_ (Tg-BK_Ca_^R207Q^) were subjected to 6hr hypoxia at 1% O_2_ followed by 12hr of reoxygenation at 21% O_2_. Cells were fixed and stained for TUNEL positive nuclei (red), and Nucleus (blue). Neonatal cardiomyocytes isolated from Tg-BK_Ca_^R207Q^ pups showed higher TUNEL positive nuclei as compared to WT-BK_Ca_ (G). Data represented as percentage TUNEL positive cells (D and G) mean ± SEM from eight of ten independent experiments. *P* values were determined by a one-tailed unpaired student’s t-test followed by a non-parametric Mann-Whitney test for comparison between the ranks; * ≤ 0.05, ** ≤ 0.001.

We have earlier shown that gain of function transgenic adult mice presented cardioprotection after ischemic preconditioning as well as after IR injury [22]. Here we isolated NCMs from wild type and Tg-BK_Ca_^R207Q^ mice[9] and subjected P3 NCMs to HR injury. After hypoxia and reoxygenation, we discovered that NCMs from Tg-BK_Ca_^R207Q^ mice showed a significant increase in TUNEL positive cells 66 ± 4% as compared to wild-type mice 17 ± 2% (**Fig. 5 E, F**, and **G**). Our data from pharmacological treatment of WT NCMs and genetically-activated transgenic mice NCMs indicate that activation of the BK_Ca_ channel increases apoptosis and blocking of the BK_Ca_ channel protects cells.

### Plasma membrane BK_Ca_ channel modulates cellular reactive oxygen species upon HR injury

Mitochondria isolated from adult guinea pig heart showed that activation of BK_Ca_ channels by NS1619 lowers reactive oxygen species (ROS) which is implicated in cardioprotection from IR injury [35-37]. We also investigated the role of plasma membrane-localized BK_Ca_ channels in regulating ROS in NCMs upon HR injury. As shown in **Fig. 6**, after hypoxia and reoxygenation, activation of BK_Ca_ channel with NS1619, intracellular ROS significantly increased compared to control and IbTx-treated cells. These results also support our findings that inhibition, and not activation of BK_Ca_ channel in NCMs or neonates is cardioprotective from IR injury.

**Figure 6:**
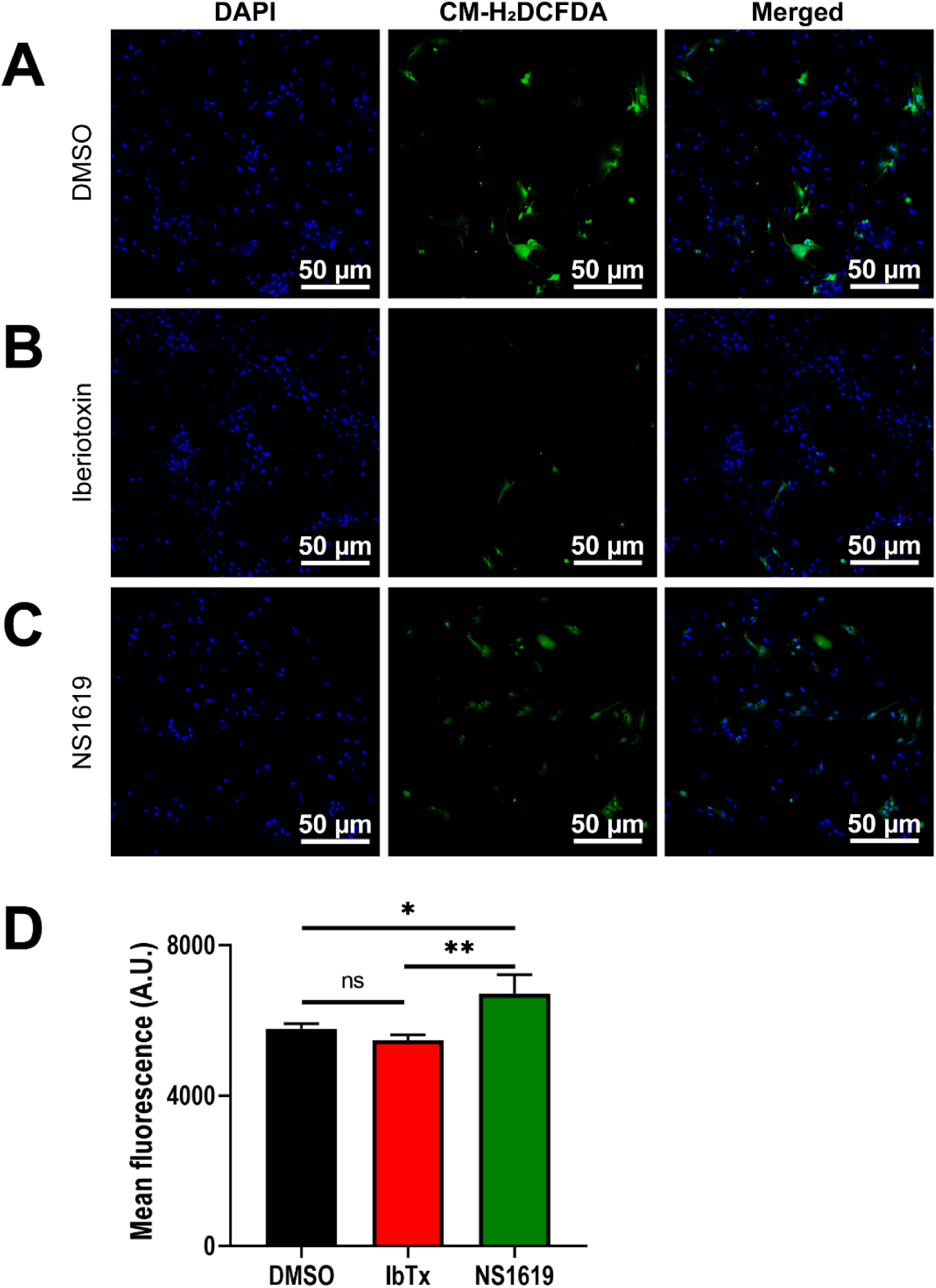
Inhibition of BK_Ca_ channel protects the neonatal cardiomyocytes from oxidative damage post hypoxia injury. Isolated rat neonatal cardiomyocytes were subjected to 6 hr hypoxia at 1% O_2_, 5 mins of post-conditioning with DMSO, Iberiotoxin (100 nM), or NS1619 (10μM) followed by 12 hr of reoxygenation. Cells were stained with CM-H_2_DCFDA (green) for 45 mins and NucBlue(tm) for nucleus (blue). A total of 100,000 cells were seeded per assay. (B and D) Cardiomyocytes post-conditioned with IbTx show less cellular ROS compared to DMSO control (A and D). Cardiomyocytes post-conditioned with NS1619 exhibited significantly higher ROS generation as compared to IbTx and DMSO control (D). Data represented as mean fluorescence unit ± SEM from eight of ten independent experiments. *P* values were determined by a one-tailed unpaired student’s t-test followed by a non-parametric Mann-Whitney test for comparison between the ranks; * ≤ 0.05, ** ≤ 0.001.

### Expression of the BK_Ca_ channel causes delayed repolarization of the plasma membrane

BK_Ca_ channel generates a large conductance for potassium in cellular membranes. Ectopic expression of the BK_Ca_ channel in murine HL-1 cardiomyocytes indicates that the introduction of BK_Ca_ shortens the action potential duration [38]. In human iPSC-derived cardiomyocytes, the BK_Ca_ channel induces irregular action potential shapes resembling very early afterdepolarizations [26]. On these lines, we performed an *in silico* simulation on cardiac action potential (AP) in presence of the BK_Ca_ channel. We assumed the open probability to 0.5 and the unitary conductance at 300 pS. The addition of the BK_Ca_ channel current creates an early voltage ‘notch’, which is noted during the depolarization phase of the AP (**Supplementary figure 1**). The peak depolarization amplitude is also reduced when compared to the native AP. Perhaps the most notable change is during the repolarization phase, which is much more rapid with the addition of the BK_Ca_ channel current. This rapid repolarization back to the resting state noticeably decreases the duration of the AP. Although these approaches indicate that induction of BK_Ca_ channel-specific current will disrupt the AP, in the absence of experimental action potential data from native cardiomyocyte BK_Ca_ channels, the precise role of the BK_Ca_ channel in cardiac AP could not be established.

Since our data indicate that a functional BK_Ca_ channel is present in NCMs, we tested whether the NCM BK_Ca_ channel plays an active role in modulating cardiac action potential. We isolated NCMs from P3 rats and seeded them on a microelectrode array for recording their activity. As shown in **Fig. 7**, NCMs are highly active and present rapid action potentials. We used two different concentrations of NS1619 (low: 10 μM and high: 20 μM) and IbTx (low: 100 nM and high: 200 nM). Low dose NS1619 or IbTx had no significant impact on APD30, APD50, APD90, and rise time of action potential, however, a high dose of NS1619 (**Fig. 7A**) impacted the NCMs AP. NCMs treated with 20 µM NS1619 showed a significantly altered AP morphology, increased APD30 (**Fig. 7C**), APD50 **(Fig. 7D**), and APD90 (**Fig. 7E**) by 4%, 11%, and 21%, respectively compared to DMSO control. Moreover, NS1619 increased the rise time (**Fig. 7F**) and APD (**Fig. 7H**) by 67% and 31%, respectively but decreased the triangulation ratio (**Fig. 7G**) by 7% when compared to DMSO. These changes in the AP morphology indicate the opening of the BK_Ca_ channel in NCMs might increase arrhythmogenic risk. Similarly, BK_Ca_ channel opening increases in the beat period (**Fig. 7K**) and field potential duration **(Fig. 7L)** by 14%, and 30% as compared to DMSO. The spike amplitude (**Fig. 7M**) showed a 30% decrease along with conduction velocity (**Fig. 7N**) which showed a 23% decrease. There were no changes observed with a high dose of IbTx (**Fig. 7B and J**) in agreement with rat and human ventricular myocyte AP’s insensitivity to IbTx [26, 39]. Adverse impact on APs and cardiomyocyte function upon opening of plasma membrane BK_Ca_ channels in NCMs implicate them in cardio-deleterious effect.

**Figure 7:**
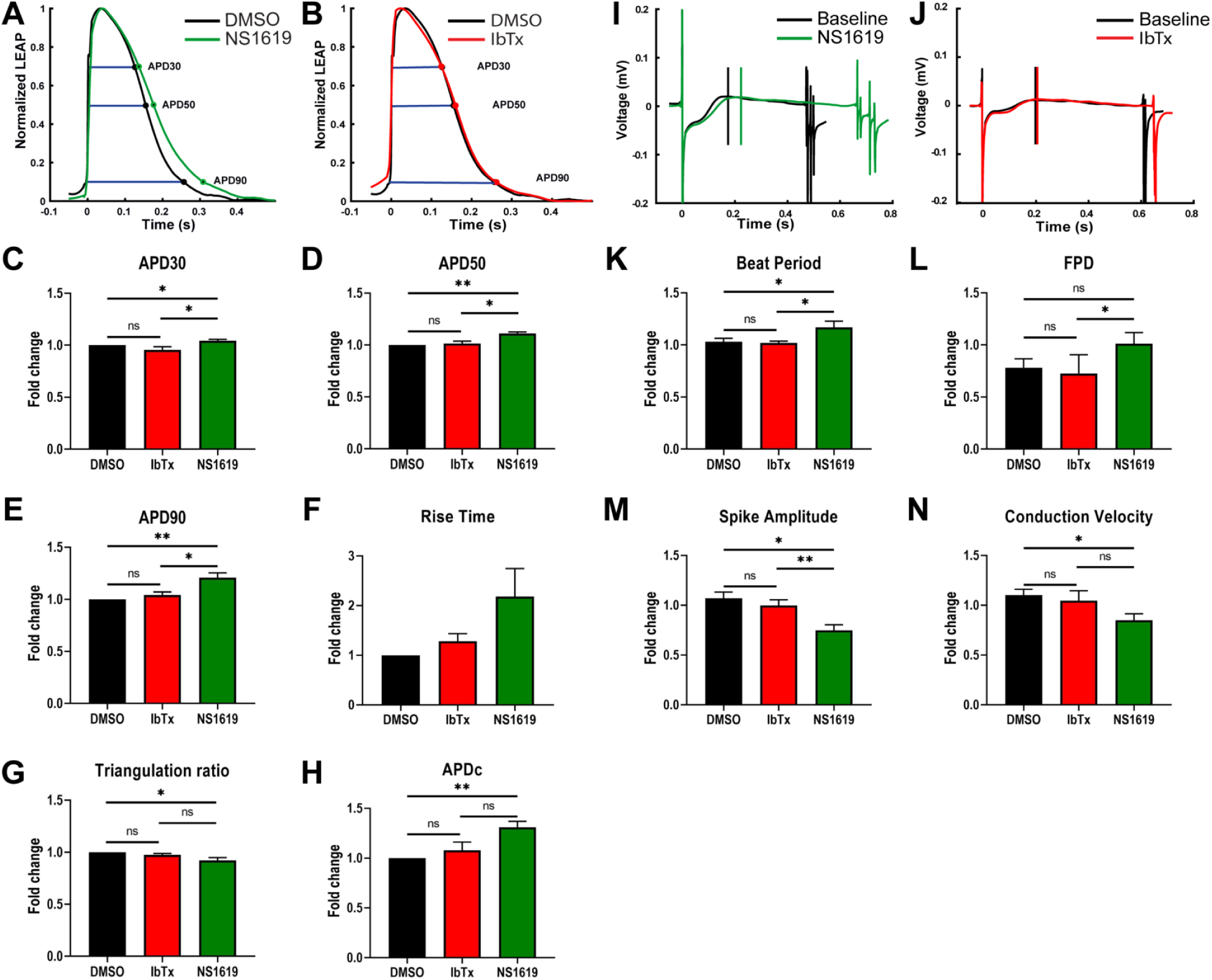
Functional analysis of activation and inhibition BK_Ca_ channel in p3 neonatal cardiomyocytes on multi-electrode array (MEA) systems. Rat neonatal cardiomyocytes plated on classic MEA 24-well plates containing electrodes were dosed with DMSO, NS1619, or IbTX. Normalized LEAP waveforms for (A; green trace) NS1619 and (B; red trace) IbTx were overlaid with DMSO control (black trace). Bar plots represent (C) APD30, (D) APD50, (E) APD90, (F) rise time (G) triangulation ratio, and (H) APDc for IbTX and NS1619 normalized to DMSO group and represented as fold change mean ± SEM. Field potential duration waveform (I and J) of neonatal cardiomyocytes post-treatment with NS1619 (I; green trace) and IbTx (J; red trace) overlaid with their respective pre-treatment baseline electrodes. Bar plots represent (K) beat period, (L) field potential duration, (M) spike amplitude, and (N) conduction velocity for DMSO, IbTx, and NS1619 were normalized to their respective pre-treatment electrodes and represented as fold change mean ± SEM. *P* values were determined by a one-tailed paired student’s t-test; * ≤ 0.05, ** ≤ 0.001.

## Discussion

Activation of the BK_Ca_ channel located in the inner mitochondrial membrane has been associated with cardioprotection and neuroprotection [29, 40]. Opening of mitoBK_Ca_ in cardiomyocytes has been implicated to increase mitochondrial K^+^ ion flux which depolarizes the mitochondrial membrane, reduces the Ca^2+^ influx and Ca^2+^ overload during injury [40]. Similarly, the opening of mitoBK_Ca_ in neurons inhibits hydrogen peroxide production which reduces ROS production and mediated neuroprotection [41]. To our knowledge, plasma membrane BK_Ca_ channel-mediated currents have not been reported in cardiomyocytes [40]. Our study for the first time indicates the presence of BK_Ca_ channels in the plasma membrane of the NCMs isolated from mice and rat hearts as well as in cardiac cells of human infant hearts. Furthermore, we have recorded K^+^ currents sensitive to IbtX and functional consequences of the activation and/or inhibition of plasma membrane BK_Ca_ channels in NCMs. Here we determined the physiological role of neonatal plasma membrane BK_Ca_ channels in cardioprotection and cardiomyocyte function.

Cardiac potassium channels are encoded by over 40 distinct genes [42]. In the heart around 10 distinct potassium channels function in tandem to tightly regulate the cardiac repolarization to ensure stable and consistent AP propagation [43]. Although coded by different genes, some of these channels have overlapping functions which result in some degree of functional redundancy and is known to contribute to repolarization reserve. Cardiac potassium channels are classified into transient outward, delayed rectifier outward, and inward rectifier currents. Canonical cardiac potassium channels play key roles during phase 1, phase 2, phase 3, and phase 4 of action potentials. More recently Ca^2+^-activated small conductance potassium channels (K_Ca_2.x) [44] and several two-pore domain potassium channels (K_2_P) [45] have been characterized in atrial repolarization. However, BK_Ca_ channels have been largely reported in cardiomyocytes’ mitochondria but not in the plasma membrane so their role in cardiac AP is not known.

Cardiac potassium channels have been associated with several cardiac diseases such as long and short QT syndromes (LQTS, SQTS), Brugada syndrome (BrS), Andersen-Tawil syndrome, and AF [46]. The occurrence of arrhythmias and ischemic heart diseases increases with cardiac development and age [47, 48]. One of the reasons attributed to the lower occurrence of cardiac electrophysiological dysfunction in the younger population could be due to differential expression of ion channels in cardiac tissues during the development. For example, in cardiac development, the expression of I_Kir_, and I_KATP_ was shown to increase from embryonic day 10 to neonate day 1 in rats, and the current density decreases after day 30 whereas I_KAch_ presented no change [49]. Ion channels play a key role in cell division and differentiation [50]. These channels need to be tightly regulated to maintain the fate and physiological role of cells. Any abnormal expression of the activity of ion channels in developing organs could have a detrimental impact.

BK_Ca_ channels are ubiquitously expressed in the plasma membrane of cardiac fibroblast [51], neurons [9, 52-56], endothelial [57, 58], and vascular smooth muscle cells [59-61]. However, in adult ventricular cardiomyocytes, a unique C-terminal splice variant of the BK_Ca_ channel targets it to the mitochondrion [13]. In the cardiovascular system, BK_Ca_ channels have been suggested to promote vascular relaxation, regulate heart rate, and protect against IR-mediated injury [13, 21-23]. In vascular smooth muscle cells, the plasma membrane-localized BK_Ca_ channel works synchronously with the ryanodine receptor and L-type calcium channel to regulate cellular contraction and relaxation [62]. Interestingly, BK_Ca_ channels have never been detected in the plasma membrane of the adult ventricular cardiomyocytes [28, 29]. On the contrary, in chick embryonic cardiac cells and hiPSC-CMs BK_Ca_ channel specific-potassium currents were recorded [26, 63], suggesting that at the early stages of cardiac development, functional BK_Ca_ channels are present in the plasma membrane of myocytes. In adult ventricular cardiomyocytes, there are no BK_Ca_ channel-specific currents [28] in the plasma membrane, which also is further corroborated by the insensitivity of IbTx on ventricular APs [26], and by immunocytochemistry [13]. In this study, for the first time in murine and rodent models, we have detected functional BK_Ca_ channels in the plasma membrane of NCMs. We have also discovered that BK_Ca_ channels localize to the plasma membrane of cardiomyocytes in human infant hearts (**Fig. 8**). The localization of BK_Ca_ channels in the plasma membrane could be transitional as cells mature and proteins reach their destined organelles. During development cardiac muscle undergoes two major changes; 1) electrical conduction and 2) energy metabolism. The presence of BK_Ca_ channels in the plasma membrane could facilitate potassium fluxes in cardiomyocytes which are essential for development. At the same time in the later part of the development, the energy demand of heart muscle increases which brings the focus to mitochondria. The presence of BK_Ca_ channels in mitochondria in adult cardiomyocytes is essential for mitochondrial structure and function[21]. Another possibility is the presence of BK_Ca_ channels in both plasma and mitochondrial membranes are essential for K^+^ homeostasis for mitochondrial function. Alternatively, it is possible that expression of BK_Ca_ channels is required to selectively remove the cells undergoing stress (hypoxia or increase in ROS) by triggering the opening of plasma membrane BK_Ca_ channels and causing apoptosis (**Fig. 6**).

**Figure 8:**
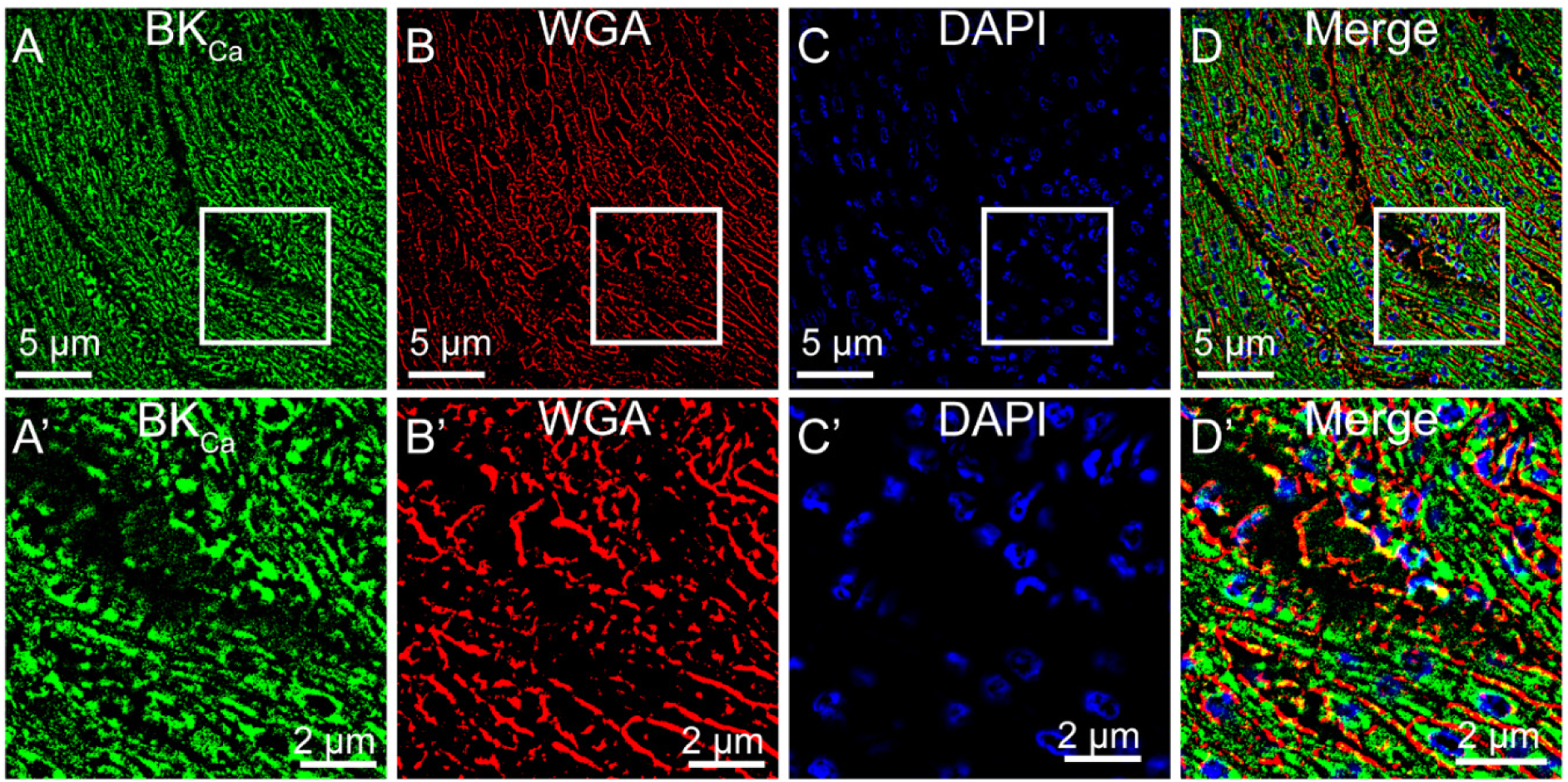
Plasma membrane localization of BK_Ca_ channel in human infant hearts. Human infant heart sections were fixed and labeled with (A) BK_Ca_ (green) (B) WGA (red), and (C) Nuclei (blue). The bottom panels (A’-D’) are shown at higher magnification with colocalization of BK_Ca_ to WGA in (D and D’).

Though no BK_Ca_ channel-specific currents were reported in the plasma membrane of adult ventricular cardiomyocytes, BK_Ca_ channels are present in the plasma membrane of smooth muscle cells. In smooth muscle cells, activation of the plasmalemma, the BK_Ca_ channel using NS1608 hyperpolarizes the membrane potential [64]. However, we have shown that blocking with IbTx implies that over ∼30% of recorded K^+^ currents originate from BK_Ca_ channels in rat and mouse NCM. In a quantitative model of a human ventricular cardiomyocyte action potential with the BK_Ca_ channel integrated into the cell membrane [26], a deep notch was shown during the depolarization phase. Our simulation of the action potential of model cardiomyocyte cells presented a similar phenotype including a notch in the depolarization phase [26]. In an overexpression model where human BK_Ca_ channels were introduced in murine HL-1 cardiac cell line [38], rapid repolarization as well as a shortened action potential were reported, which is consistent with our simulation. These *in silico* and experimental data indicate an opening of the BK_Ca_ channel allows rapid efflux of K^+^ in response to voltage and [Ca^2+^]_i_, which aids in rapid repolarization and thus shortening of the AP. However, little is known about the functional consequences of uniquely localized native plasma membrane BK_Ca_ channel in neonatal cardiomyocytes.

Due to the distinct localization of the native BK_Ca_ channel in the plasma membrane, we studied the effect of its activation and inhibition on the electrical activation of NCMs. In HL-1 cardiac cells, overexpression of the BK_Ca_ channel affects repolarization K^+^ currents and shortening of action potential duration without depolarizing the membrane or impacting intracellular Ca^2+^ influx [38]. In agreement, our data indicated that activation of endogenous BK_Ca_ channel present on NCM plasma membrane shows no change in the membrane depolarization. However, activation of endogenous BK_Ca_ channel delays the repolarization and severely prolongs the duration of the action potential. On the contrary, inhibiting the BK_Ca_ channel activity using IbTx, did not affect the repolarization or duration of the action potential. Since these adverse consequences were observed on the opening of BK_Ca_ channels, our results suggest that under physiological conditions plasma membrane BK_Ca_ channels in NCM are not active.

In adult experimental models, activation of BK_Ca_ channels by NS1619 and NS11021 protected the hearts against IR injury [24, 32, 65-71] and inhibition of BK_Ca_ channels by IbTx or paxilline increased myocardial infarction or reversed cardioprotective effects caused by the opening of BK_Ca_ channels [23, 32, 33, 69, 72, 73]. Pharmacological data were inadvertently supported by genetic models comprising global [13], cardiomyocyte-specific [23], or gain-of-function transgenic mice for BK_Ca_ channels [22]. The cardioprotective mechanism mediated by the BK_Ca_ channel is associated with an increase in mitochondrial ROS generation and Ca^2+^ overload which in turn triggers the formation and opening of mitochondrial permeability transition pore [21]. These findings are in agreement with the notion that stored potassium in intracellular organelles such as mitochondria protects cells from apoptosis [74, 75].

In our study, we discovered that the opening of BK_Ca_ channels in an organ or cellular level causes apoptosis and increases myocardial infarction. The contrasting outcome of activation of BK_Ca_ channels in neonates as opposed to adults could be attributed to the plasma membrane localization and opening of BK_Ca_ channels. Our data indicate that the opening of BK_Ca_ channels at the plasma membrane in neonates will cause membrane depolarization which in turn will trigger cell death. Under physiological conditions, BK_Ca_ channels are not anticipated to open in neonates which will not cause any adverse effects on cardiac cells. However, under pathological conditions, such as dilated cardiomyopathy in children, there is evidence of increased sensitivity of myocytes to Ca^2+^, and pediatric cardiomyocytes have decreased cooperativity when compared with adult cardiomyocytes [76]. The increase in cellular Ca^2+^ could trigger the opening of the plasma membrane BK_Ca_ channel which will immediately cause depolarization and heart failure. This could be one of the key reasons why the use of proven adult heart failure pharmacological interventions in pediatric or children with dilated cardiomyopathy do not present the same beneficial outcome as adults. Therefore, it is important to take the presence of the BK_Ca_ channel into account before selecting the appropriate pharmacological therapies in the pediatric population.

## Statements and Declarations

None

## Availability of data and materials

All data and materials are available on reasonable request to the corresponding author.

## Acknowledgments

We are grateful to Dr. Jianli Bi in the Heart Center Biorepository at Nationwide Children’s Hospital, Columbus, OH, for providing technical help with human infant myocardial sections. The authors thank Ms. Manasvini Kumaraswamy and Ms. Yashika Kuchallapali for their help with blinded analysis. We would also like to thank Prof. Robin Shaw (University of Utah) for a consultation about *in-silico* analysis and Prof. Andrea Meredith (University of Maryland) for providing us with BK_Ca_-TG mice.

## Author contributions

SS: designed and performed experiments, and assisted with interpretation of results. KS: performed electrophysiology of NCMs and analyzed data. DP: assisted in cell biology and imaging. DS: assisted in MEA experiments. AL: performed electrophysiology of NCMs. IH: whole-cell patch-clamp on P6 NCMs. AC: performed in silico experiments. UM: assisted in human heart processing. ARK: assisted in *in silico* data analysis. SGR: assisted in cell biology data analysis. MK: assisted in MEA data analysis. VG: assisted with acquisition and analysis of human hearts. HS: performed ex-vivo IR injury assay, and assisted with designing and analysis of experiments, and performed interpretation of results and manuscript preparation.

## Funding

This work was supported by grants from the National Institutes of Health (NIH) R01-HL133050 (HS), American Heart Association (AHA) Career Development Award (20CDA35310714, DP) and AHA Scientist Development grant (11SDG7230059, HS) and W. W. Smith Charitable Trust (HS).

## Ethics declarations

### Conflict of interest

The authors declare that they have no conflict of interest.

### Ethical approval

All animal handling and laboratory procedures were in accordance with the approved protocols (2018A00000095) of the Institutional Animal Care and Use Committee at the Ohio State University conforming to the NIH Guide for the Care and Use of Laboratory Animals (8th edition, 2011).

## Notes

### Competing Interest Statement

The authors have declared no competing interest.

