## Supplementary information for "Inhibition of BK_Ca_ channels protects neonatal hearts against myocardial ischemia and reperfusion injury"

**Supplementary Figure 1**

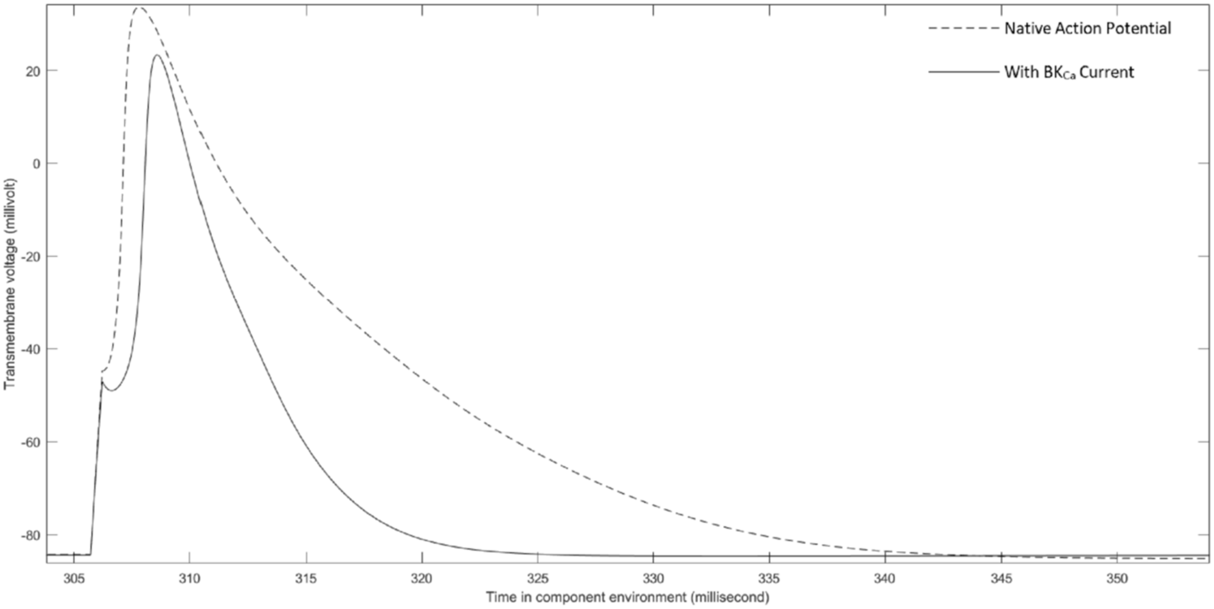


**Supplementary Figure 1. Alteration of adult murine ventricular cardiomyocyte AP with the integration of $BK_{Ca}$ channel into the cell membrane**. As compared to the native action potential (dotted line), integration of the $BK_{Ca}$ current into the murine ventricular AP (solid line) causes an early notch in the action potential, decreased maximum depolarization amplitude, and more rapid repolarization which decreases the duration of the action potential.

**Supplementary Table 1. Primers for qPCR.**

| Splice variant | Primer | Sequence | Size (bp) |
|---|---|---|---|
| Total | Forward primer | 5’ CCATTAAGTCGGGCTGATTTAAG 3′ | 187 |
| | Reverse primer | 5′ CCTTGGGAATTAGCCTGCAAGA 3′ | |
| $BK_{Ca}$-DEC | Forward primer | 5′ GGTTTACAGATGAGCCGGATA 3′ | 134 |
| | Reverse primer | 5′ CATCTTCAACTTCTCTGATTGG 3′ | |
| GAPDH | Forward primer | 5’ ACAGCAACAGGGTGGTGGA 3’ | 117 |
| | Reverse primer | 5’ TTTGAGGGTGCAGCGAACTT 3’ | |
